# Development of an Accelerated Cellular Model for Alzheimer’s Disease

**DOI:** 10.1101/2023.05.09.539465

**Authors:** Huijing Xue, Sylvester Gate, Emma Gentry, Wolfgang Losert, Kan Cao

## Abstract

Alzheimer’s Disease (AD) is a leading cause of dementia characterized by amyloid plaques and neurofibrillary tangles, and its pathogenesis remains unclear. Current cellular models for AD often require several months to exhibit phenotypic features due to the lack of an aging environment in vitro. Lamin A is a key component of the nuclear lamina. And progerin, a truncated protein resulting from specific lamin A mutations, causes Hutchinson-Gilford Progeria Syndrome (HGPS), a disease that prematurely ages individuals. Studies have reported that lamin A expression is induced in the brains of AD patients, and overlapping cellular phenotypes have been observed between HGPS and AD cells. In this study, we investigated the effects of exogenous progerin expression on neural progenitor cells carrying familial AD mutations (FAD). Within three to four weeks of differentiation, these cells exhibited robust AD phenotypes, including increased tau phosphorylation, amyloid plaque accumulation, and an elevated Aβ42 to Aβ40 ratio. Additionally, progerin expression significantly increased AD cellular phenotypes such as cell death and cell cycle re-entry. Our results suggest that progerin expression could be used to create an accelerated model for AD development and drug screening.

**Significance Statement:** Alzheimer’s Disease (AD) contributes to most dementia, while its mechanism is still under investigation. One of the challenges for studying AD is the model issue, including the genetic divergence of animals and human, and the rejuvenation of induced pluripotent stem cells (iPSCs). Progerin is a mutant lamin A found in the accelerated aging disease progeria. There are a lot of molecular similarities between Alzheimer’s Disease (AD) and progeria. Here, we developed an accelerated 2D/3D cell model system for AD by ectopically expressing progerin in a previously characterized AD cell model carrying familial AD (FAD) mutations. Our study showed that progerin addition could accelerate AD phenotypical progression, including tau phosphorylation and formation of β-amyloid plaques.

## Introduction

The nuclear lamina, located under the inner nuclear membrane, provides structural support to the nucleus and interacts with chromatin and inner nuclear membrane proteins (1). It mainly consists of two types, A-type lamins (lamin A and lamin C) and B-type lamins (lamin B1 and lamin B2). Lamin A and lamin C are the alternatively spliced products of the *LMNA* gene (2), and lamin B1 and lamin B2 are encoded by *LMNB1* (3) and *LMNB2* (4), respectively. Mutations in *LMNA* lead to a wide range of diverse diseases called laminopathies. In particular, Hutchinson-Gilford Progeria Syndrome (HGPS), a devastating premature aging disease, is mainly caused by a point mutation in the exon 11 of the *LMNA* gene, G608G (GGC > GGT), resulting in a cryptic splicing event and generates a truncated product called progerin (5). The truncated part contains a ZMPSTE24 cleavage site, which is responsible for the cleavage of the farnesylated tail in the prelamin A (5). Thus, progerin is permanently farnesylated and becomes attached to the inner nuclear membrane (INM) (6). The accumulation of progerin inside the nucleus causes misshaped nuclear morphology, lamin B1 downregulation, chromatin relaxation, and apoptosis (7), which overlap with the phenotypes in the Alzheimer’s Disease (AD) model (8).

AD is a neurodegenerative disease, and one of the most common causes of dementia (9). It can be subtyped into two categories. One is familial AD (FAD), which begins before age 65, and the other is sporadic AD (SAD), which usually begins after age 65 (10). Two hallmarks of AD are senile plaques made up of β-amyloid (Aβ) and neurofibrillary tangles (NFTs) made up of tau protein (11). Both genetic and environmental risk factors can contribute to the formation of plaques and fibrillary tangles, with aging being one of the greatest risk factors (12). However, the specific mechanism underlying how these protein aggregations form remains unclear. One leading hypothesis is Aβ cascade hypothesis. Aβ is derived from the Amyloid Precursor Protein (APP), and it has different isoforms, Aβ40, Aβ42, and Aβ43, which are the products of heterogeneous γ-secretase cleavage (13). Starting from monomers, Aβ aggregates into oligomers and fibrils, causing neurotoxicity and leading to neuronal cell death and neurodegeneration (14). Aβ42 is highly self-aggregating and, therefore, potentially promotes brain Aβ deposits (15)’(16); thus, Aβ42 to Aβ40 ratio is considered a sensitive diagnostic marker(17). Another main hypothesis for AD mechanism involves tau phosphorylation. Tau belongs to Microtubule-Associated Proteins (MAPs) and acts as a microtubule stabilizer in the axon (18). Tau protein is hyperphosphorylated in the brains of AD patients, and these abnormally phosphorylated proteins form paired helical filaments (PHF), and further form NFTs (19). There are different sites of tau phosphorylation, including Tyr18, Ser199, Ser202/Thr205, Thr231, Ser262, Ser396, Ser422 (20). Whether Aβ cascade and tau phosphorylation are causal or synergistic is still under investigation (21–23). Both of these pathways are reported to be involved in AD events, including cell cycle re-entry, cell apoptosis, and oxidative stress(24–26).

Several studies have suggested the potential involvement of the nuclear lamina in AD. Researchers reported the changes in nuclear morphology in various AD models (27, 28). Despite the low expression level of lamin A in the brain, one group observed that both transcriptional and translational levels of lamin A were significantly increased in the hippocampus in late-stage AD (29). Another group also observed a significant increase of lamin A expression and reinforced perinuclear Lamin B2 in hippocampal neurons through AD progression (30). Besides AD, a strong upregulation of lamin A can be detected in the brain of Alexander disease (AxD) patients, and it is associated with increasing brain stiffness (31). Moreover, overexpression of A-type lamins, including the mutant progerin, is known to negatively impact B-type lamins’ expression (32), which can induce heterochromatin relaxation in AD (28). These studies suggest that lamin A accumulation may play a role in neurodegeneration.

One of the biggest challenges for studying AD is accurately and efficiently modeling the disease. Considering the significant sequence differences in Aβ and tau between mice and humans, human tissue and cells could provide more accurate information (33, 34). Because of the lack of high-quality human post-mortem tissues, cell culture models have become a popular choice. Most AD cell culture models are derived from human induced pluripotent stem cells (iPSCs) (35). However, iPSC-derived neurons can be rejuvenated after reprogramming (36). This poses a concern as while AD is an age-related disease, which could explain why many current iPSC-derived models are time-consuming and fail to recapitulate the amyloid plaque and tau tangle simultaneously. The appearance of AD characteristic features in primary cell culture often requires several months (37–39), and the time for detecting these phenotypes can be unpredictable, making data reproducibility a challenge. Thus, developing an efficient AD cellular model is urgent. Considering the overlapping cellular phenotypes between HGPS and AD and the induced expression of Lamin A in AD brains (8), we propose to study whether the ectopic expression of progerin could be a useful strategy to promote an aging environment in the *in vitro* cellular models for better and faster modeling of AD.

To test this idea, we chose to modify one of the leading cellular models for AD, which requires several months to develop and display AD phenotypes documented by various publications (38, 40). We found that progerin expression could accelerate AD phenotype exhibition from 8-16 weeks to 3-4 weeks, providing a faster, cheaper, and more predictable platform for modeling AD.

## Results

### Different nuclear lamins are regulated differently during neural differentiation

We first characterized the expression of various lamins during the neural differentiation of ReN cells, a commercial human neural progenitor cell (NPC) line (Fig S1). By retracting growth factors from the medium, ReN cells generated neurite morphology (Fig S1a). To validate the cell types, we detected the cells with neuronal marker MAP2 and astrocytes marker GFAP by immunofluorescence staining. As expected, both markers were positively stained after two-week differentiation (Fig S1b). To further determine if these cells were functional, we stained the differentiated cells with a Ca2+ indicator, Fluo-4. By measuring the fluorescence change, the Ca2+ transient could be detected and therefore reflect the action potentials (41). Cells at day 14 could generate Ca2+ transients regularly when the medium was replaced by Tyrode’s solution supplemented with Na+ (Fig S1c). This result suggested that ReN cells could generate action potentials after differentiation.

To determine the expression pattern of nuclear lamins in neurons, we probed both transcriptional and translational expression of nuclear lamins during differentiation (Fig 1). We found that at the mRNA level, both lamin A and lamin B1 were decreased, while lamin C did not show a significant change (Fig 1a). However, at the protein level, lamin A and lamin C were downregulated, while lamin B1’s protein level was relatively stable (Fig 1c,d). These results suggested that different nuclear lamins have different stability and different regulation mechanisms. While lamin A exists in the neural progenitor cells, it is diminished after the differentiation, which is consistent with previous findings (42).

**Figure 1.**
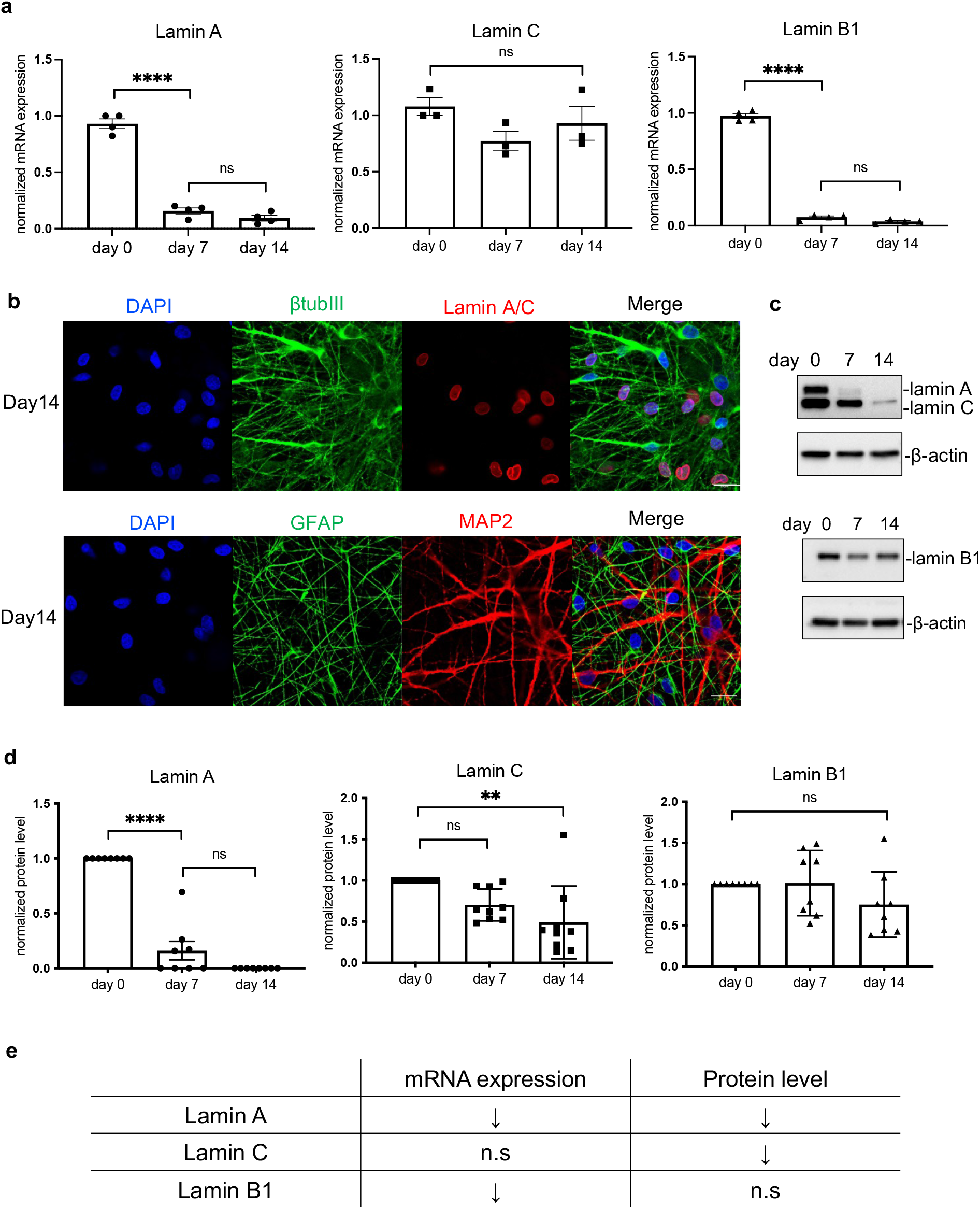
The expression of nuclear lamins mRNA and protein during the differentiation. (a)The quantification of mRNA relative expression of lamin A, lamin C and lamin B1 at different time points (day 0, day 7, day 14). Both lamin A and lamin B were significantly decreased while there was no significant change in lamin C mRNA. Results were generated from more than three biological replicates. Asterisks indicate statistical difference as follows: ns, not significant; *p < 0.05; **p < 0.01; ***p < 0.001; ****p < 0.0001. (b)Immunofluorescence staining of neuronal markers and lamin A/C at day 14. β-tubulin III and MAP2 are neuronal markers and GFAP is astrocytes markers. Lamin A/C staining is also positive in ReN cells after two-week differentiation. (Scale bar: 20um) (c)(d) Western blot results of lamin A, lamin C and lamin B1 protein levels at different time points (day0, day7, day14). Both lamin A and lamin C were downregulated while lamin B1 was relatively stable. Results were generated from more than three biological replicates. Asterisks indicate statistical difference as follows: ns, not significant; *p < 0.05; **p < 0.01; ***p < 0.001; ****p < 0.0001. (e)Summary of the mRNA and protein expression pattern of nuclear lamins during NPC differentiation.

### Overexpression of lamin A and progerin results in neural death and cell cycle re-entry

Healthy mature neural cells prefer expressing lamin C rather than lamin A (42). However, abnormal lamin A accumulation has been observed in patients’ hippocampus through the different stages of AD (29, 30). To probe the potential role of A-type lamins in neurodegeneration, we overexpressed lamin A and progerin, respectively, in differentiating ReN cells to check their effects. Lentivirus constructs were used to express lamin A and progerin in ReN cells (Fig S2a), as previously described (43). Both lamin A and progerin were tagged with EGFP (44). Flow cytometry analysis indicated that transduction efficiencies were 71.1% and 64.8% in lamin A and progerin, respectively (Fig S2b, c). After transduction, ReN cells underwent the differentiation process. We checked the exogenous protein expression during the differentiation. By quantifying the protein level with Western blot (Fig S2d), and found that exogenous lamin A and progerin were still abundant after 2-week differentiation but were decreased after 4 weeks.

Oxidative stress increases during aging and it is extensively reported that oxidative imbalance plays a critical role in AD (45). To check the oxidative stress in ReN cells, we measured Reactive Oxygen Species (ROS) level. We observed increased ROS levels after lamin A expression, and progerin overexpression could significantly intensify this effect in 2 weeks (Fig 2a). Cell death is another widely accepted feature of AD (46), and we performed cell death flow cytometry to check this phenotype. After two weeks of differentiation, we observed a slight increase in the proportion of early cell death events in cells expressing lamin A compared to non-transduced cells (Fig 2b). Moreover, progerin-transduced cells showed a significant increase in cell death compared to non-transduced cells at the same time point.

**Figure 2.**
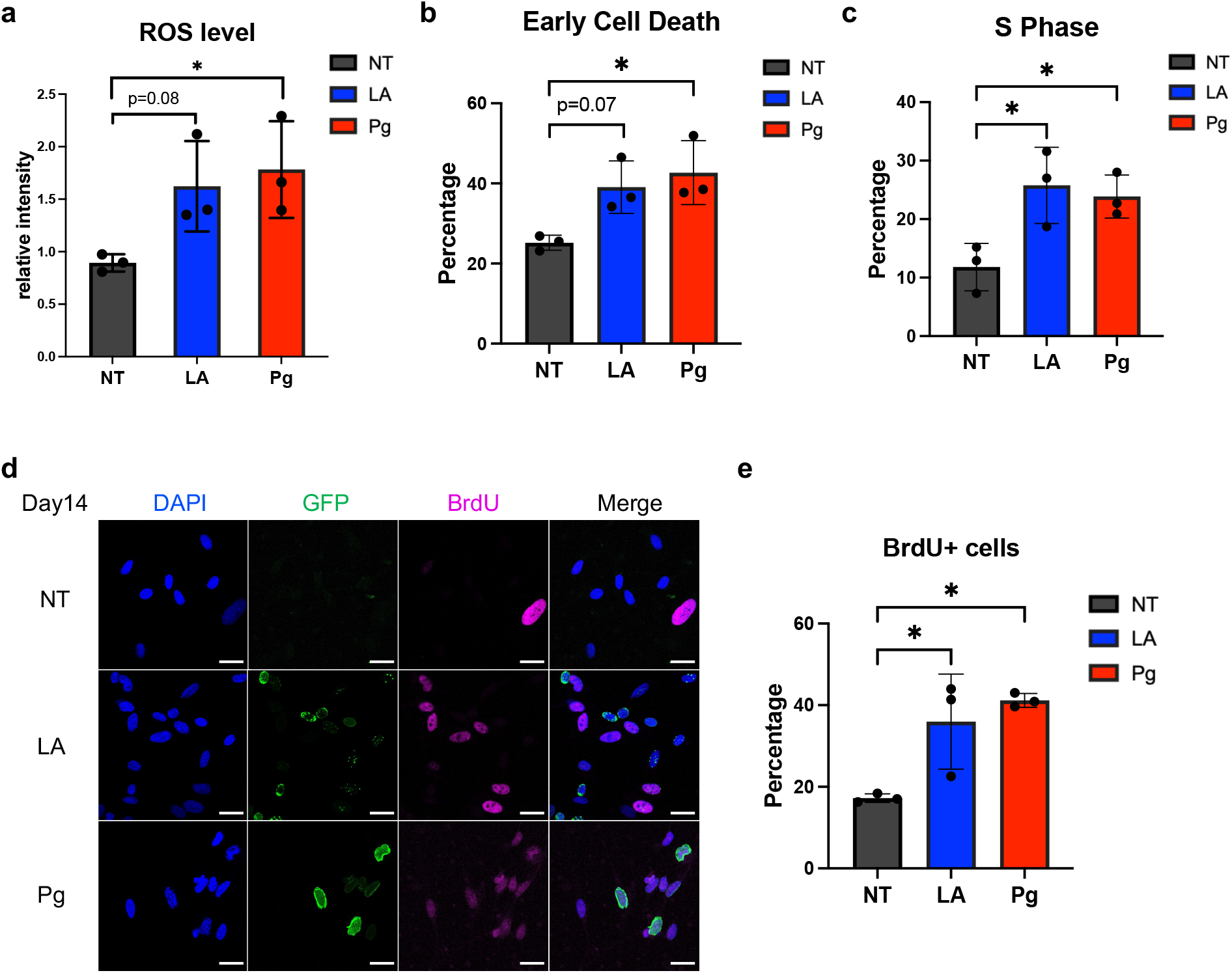
The effects of the overexpression of lamin A and progerin in neural cells. (a)ROS level after two-week lamin A and progerin overexpression. ROS level was significantly increased with progerin expression. Results were generated from three biological replicates. *p < 0.05. (b)The quantification of cell death assay. The percentage of early cell death was significantly increased with progerin expression after two-week differentiation. Results were generated from three biological replicates. *p < 0.05. (c)The quantification of cell cycle assay. The percentage of S-phase cells was increased after two-week lamin A- or progerin-transduction. Results were generated from three biological replicates. *p < 0.05. (d)(e) Immunofluorescence staining of BrdU in differentiated cells. Blue indicated DAPI signal, green indicated GFP-tagged lamin A or progerin, pink indicated BrdU signal. The percentage of BrdU positive cells was increased after lamin A- or progerin-transduction. Results were generated from three biological replicates. *p < 0.05. (Scale bar: 20um)

Although neurons are known to quit the cell cycle and stay in the quiescent stage (G0), cell cycle re-entry is considered an early event in neurodegeneration and is related to cell death (47, 48). A significantly higher percentage of S-phase cells was detected after 2-week lamin A-transduction, compared to the cells without any transductions (Fig 2c), which indicated increased cell cycle re-activation. We also stained the cells with BrdU to further validate the S-phase cells and confirmed that the percentage of S-phase cells was increased after 2 weeks (Fig 2d,e). In summary, the overexpression of lamin A led to heightened oxidative stress, cell death, and cell cycle re-entry, while progerin further exacerbated these characteristics.

### The combination of progerin and FAD mutations accelerated AD hallmark phenotypes

As stated previously, the age clock is reset in iPSC-derived neurons even if the cells are from old donors (36). In addition, *in vitro* cell culture often positively selects the more proliferative (i.e., younger and healthier) cells. Therefore, modeling an age-associated disease *in vitro* is tricky. Since we found that ectopic lamin A- or progerin-expression could induce age-associated phenotypes ReN cells after 2-week differentiation (Fig 2), we asked if lamin A- or progerin-expression could accelerate neurodegeneration in ReN cells with FAD mutations.

To test this idea, we adapted a well-characterized AD model (38), which is the first cell culture model recapitulating both Aβ plaques and tau aggregations. The plasmid containing APP with both the K670N/M671L (Swedish) and V717I (London) mutations (APPSL) and PSEN1 with the Δ9 mutation (PSEN1(Δ9)) was utilized to introduce FAD mutants (Fig S3a), as described (38). We transduced the ReN cells with lentivirus containing <u>m</u>cherry-tagged <u>A</u>PPSL and <u>P</u>SEN1(Δ9) mutations (mAP) before the differentiation and used mcherry-only lentivirus as the control group. For 2D culture, cells were seeded on the Matrigel-coated plates. For 3D culture, the cell suspension was mixed with Matrigel with a certain ratio and then plated on the chamber slides. After 3 days, both APP mRNA and PSEN1 mRNA were increased after mAP transduction (Fig S3b). After a 2-week differentiation period, cells were transduced with either GFP-lamin A (LA) or GFP-progerin (Pg). Two days after transduction, both lamin A and progerin expression were robust. However, after 2 weeks, while lamin A expression remained strong, progerin expression became very weak.

Tau phosphorylation is an important hallmark of AD (18). We used Western Blot to check the protein level of total tau and phosphorylated tau weekly. It is reported that the phosphorylation of Thr231 tau is an early event in AD among different phosphorylation sites (49). It is estimated that ptau at Thr231 contributes about 26% of the overall inhibition of tau binding to microtubules (50). Thus, an antibody that recognizes Thr231 phosphorylation site was used to detect tau phosphorylation. Although the total tau level was not significantly changed, ptau at Thr231 was slightly increased after lamin A expression compared to the cells with FAD mutations only, and progerin expression further upregulated tau phosphorylation significantly at week 4 (Fig 3b, c).

**Figure 3.**
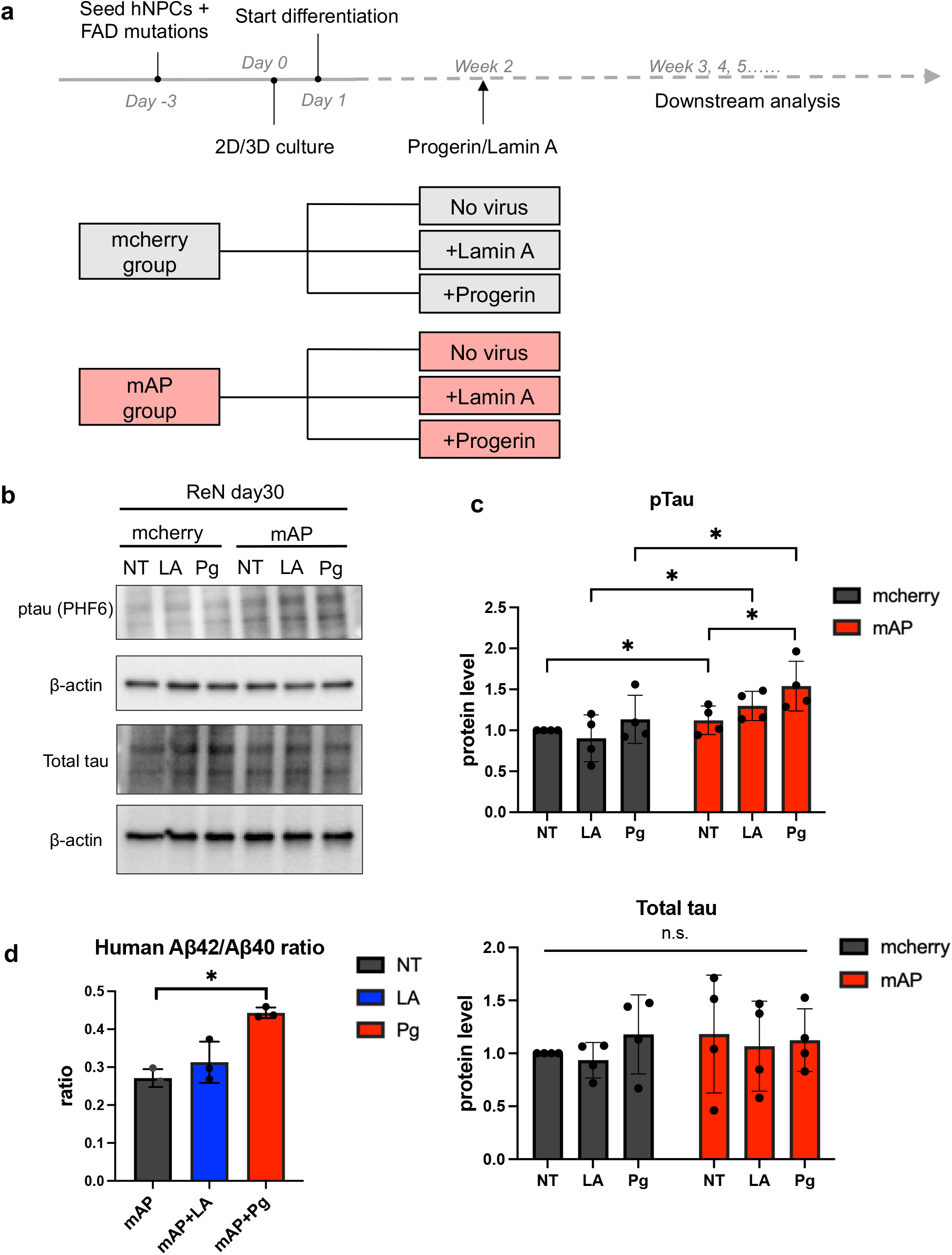
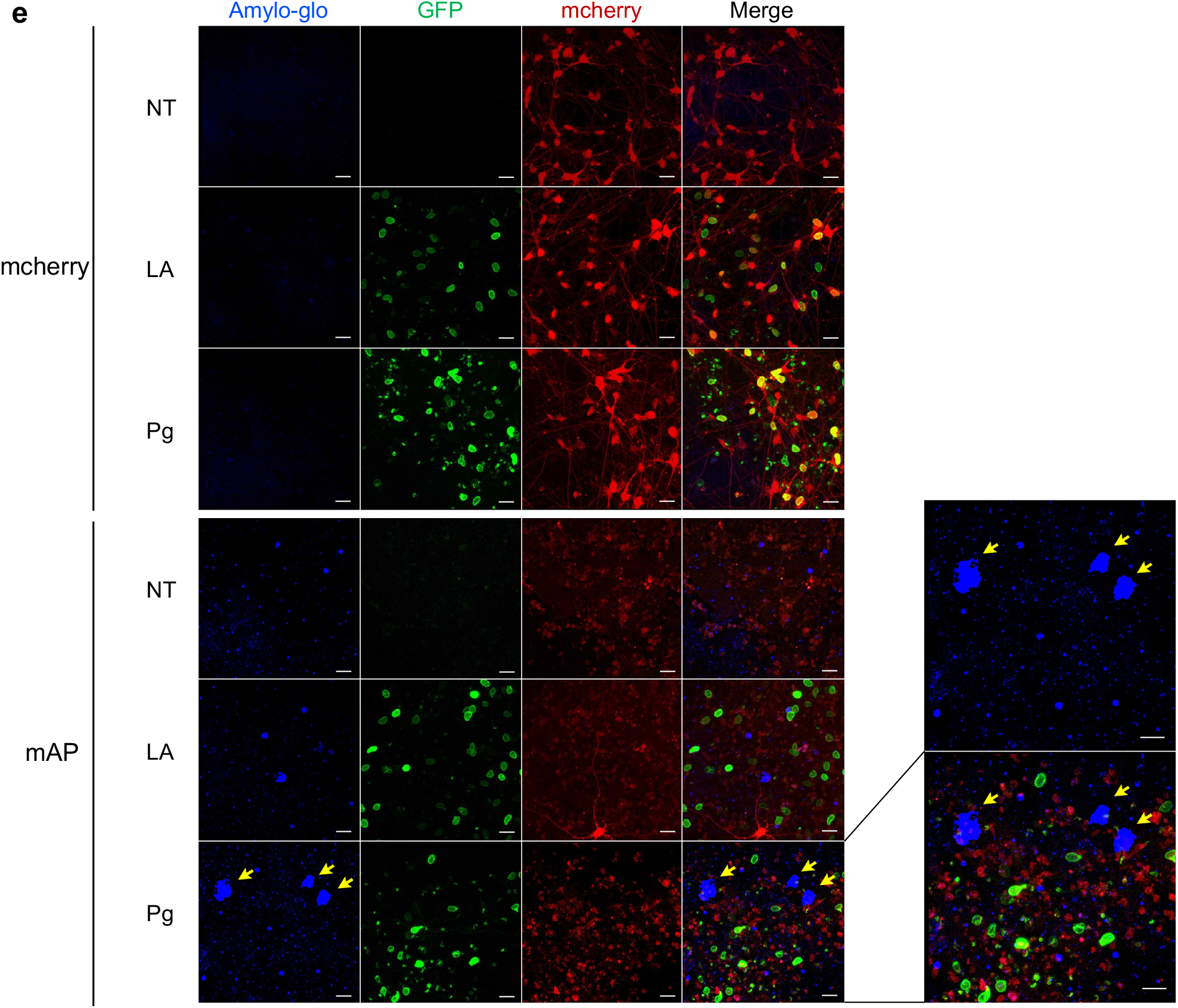
Accelerated AD protein features after FAD mutations and progerin co-transduction. (a) Experimental timeline. ReN cells were transduced with the lentivirus containing FAD mutation constructs before the differentiation. Cells were seeded on Matrigel-coated plates for 2D culture while cell suspension was mixed with Matrigel in 3D culture. After two-week differentiation, lamin A or progerin expression were transduced in the cells and followed by downstream analyses. Samples were as listed. (b)(c) Protein level of total tau and phosphorylated tau after 4 weeks. The total tau level was not significantly changed. Phosphorylated tau was increased after lamin A expression, and progerin expression further upregulated tau phosphorylation significantly. Results were generated from four biological replicates. n.s., not significant; *p < 0.05. (d)Aβ42/ Aβ40 ratio after 3-week differentiation. Within the mAP group, Aβ42/ Aβ40 ratio was slightly increased after lamin A expression and significantly increased after progerin expression. Results were generated from three biological replicates. *p < 0.05. (e)Aβ aggregation staining with Amylo-glo after 4-week differentiation. Blue indicated the Amylo-glo staining, green indicated the GFP-tagged lamin A or progerin, red indicated mcherry or mcherry-tagged APP and PSEN1. Yellow arrows indicated the Aβ fibrils. (Scale bar: 20um)

Amyloid is another crucial pathology marker for AD (51). A higher Aβ42/ Aβ40 ratio usually indicates more neurotoxicity, and it could be used as a more sensitive marker (17). We used ELISA to detect the Aβ40 concentration, Aβ42 concentration, and Aβ42/ Aβ40 ratio in the culture medium after 3-week culture. Aβ40 level was upregulated in the mAP group after 3 weeks (Fig S3d). Aβ42 was barely detectable in the mcherry control group but was again upregulated in the mAP group after 3 weeks (Fig S3d). In the mAP group, the Aβ42/Aβ40 ratio showed a slight increase following the introduction of lamin A compared to cells carrying FAD mutants alone, and progerin expression significantly raised the Aβ42/Aβ40 ratio in comparison to cells with FAD mutants alone (Figure 3d).

Next, we checked for any Aβ aggregation formations weekly in both 2D and 3D culture. In the 2D cell culture, we checked the cells with an Aβ oligomer antibody. In mcherry group, Aβ oligomers were barely detected. In the mAP group, cells expressing both FAD mutations and progerin displayed stronger Aβ oligomer staining compared to cells expressing only FAD mutations after 3 weeks of culture (Fig S3e). Aβ usually diffuses into media in 2D cell culture and therefore it is difficult to detect fibril formation. To further visualize the aggregation, a 3D cell culture with Matrigel was adapted, and Amylo-glo was used to detect the Aβ fibrils weekly. Amylo-glo signals were first detected in the mAP group after four weeks of culture, whereas the signal remained negative in the mcherry control group (Fig 3e). Notably, we observed larger Aβ fibrils in the mAP cells expressing both progerin and FAD mutations after four weeks, as compared to the cells expressing FAD mutations alone (Fig 3e). These results indicated that ReN cells with the combination of progerin and FAD mutants displayed accelerated disease phenotypes after only 3-4 weeks in both 2D and 3D cell culture, including tau phosphorylation and formation of β-amyloid.

### The combination of progerin and FAD mutations leads to increasing cell cycle re-entry and increasing cell death

After checking the AD pathological hallmarks, we investigated how cell cycle reactivation changed with the addition of lamin A or progerin. Overall, progerin addition resulted in more cell cycle re-entry events in 4 weeks (Fig 4a). Specifically, a mild increasing percentage of S-phase cells were observed, comparing cells with FAD mutants alone to those with mcherry control plasmid. Within the mcherry group, ectopic expression of lamin A slightly increased the percentage of S-phase cells, and progerin could lead to a more drastic increase in S-phase cells after 4 weeks, compared to the cells with mcherry signal alone. Cells with FAD mutants exhibited a significantly higher percentage of S-phase cells after the ectopic expression of either lamin A or progerin. We further checked cell cycle re-entry with BrdU staining (Fig 4b, S4). The combination of progerin and FAD mutations induced more BrdU+ cells than FAD mutations alone after 4-week culture. The same trend could be observed for the combination of lamin A and FAD mutations (Fig 4b, S4).

**Figure 4.**
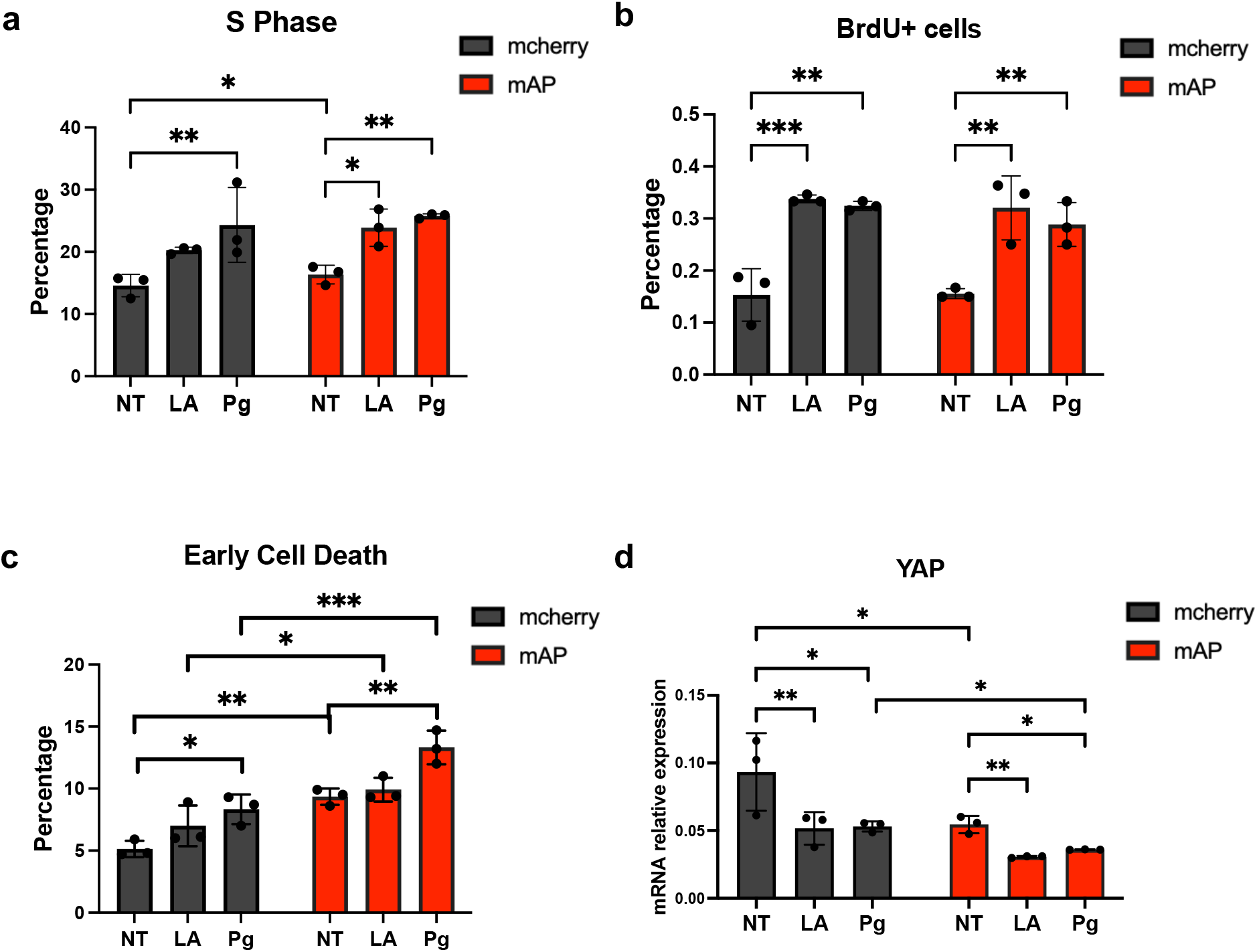
Increased cell cycle re-entry and cell death after FAD mutations and progerin co-transduction after 4 weeks. (a)The quantification of cell cycle assay. Within both mcherry control group and mAP group, S phase cells were increased after lamin A- and progerin-transduction. Comparing mAP cells to mcherry control cells, S phase percentage was increased as well. Results were generated from three biological replicates. n.s., not significant; *p < 0.05; **p < 0.01; ***p < 0.001; ****p < 0.0001. (b)The quantification of BrdU positive cells. An increased percentage of BrdU+ cells after lamin A- and progerin-transduction was observed in both mcherry control group and mAP group. Results were generated from three biological replicates. n.s., not significant; *p < 0.05; **p < 0.01; ***p < 0.001; ****p < 0.0001. (c)The quantification of cell death flow cytometry. Cell death was increased comparing mAP group to mcherry control group. Within each group, progerin expression significantly induced more cell death. Results were generated from three biological replicates. n.s., not significant; *p < 0.05; **p < 0.01; ***p < 0.001; ****p < 0.0001. (d)The quantification of mRNA relative expression of YAP. YAP mRNA was downregulated in mAP ReN cells and progerin expression could further decrease YAP expression. Results were generated from three biological replicates. n.s., not significant; *p < 0.05; **p < 0.01; ***p < 0.001; ****p < 0.0001.

It is reported that increased p16 and cdk4/6 are associated with cell cycle dysregulation in AD(52, 53). Thus, the translational expression of these cell cycle regulators was investigated. In general, the intervention of progerin gave rise to the significantly elevated mRNA level of p16 and cdk4/6 in both mcherry control group and mAP group in 4 weeks (Fig S5). This suggested that p16, cdk4/6 were involved in progerin-induced cell cycle reactivation. Ectopic expression of lamin A had the same trend (Fig S5). Although the transcriptional expressions of p16 and cdk4 were indistinguishable between cells with FAD mutants and cells with mcherry control plasmid (Fig S5a,b), cdk6 mRNA was significantly higher in cells with FAD mutants (Fig S5c).

Since cell cycle re-entry has been associated with cell death in neural cells (54), we performed cell death flow cytometry. Results showed that progerin addition significantly induced more cell death events in both mcherry control group and mAP group in 4 weeks (Fig 4c). Comparing each parallel sample in mAP group to the sample in mcherry control group, significantly increased cell death was detected as well (Fig 4c). These results indicated that progerin and FAD mutants could have a synergetic effect in cell death.

Cell death is a critical event in AD progression (46) and we would like to check the potential reasons in our system. Yes-associated protein (YAP) is a critical factor of the Hippo signaling pathway, which responds to changes in cell mechanics (55). Several studies suggested YAP was downregulated in AD brains (56) and it could be a critical regulator for cell death in AD (57). Here we checked the translational expression of YAP. YAP mRNA was significantly downregulated in the cells with FAD mutants after 4 weeks, compared to the mcherry control group (Fig 4d). The combination of progerin and FAD mutants exhibited a further reduction of YAP, compared to the cells with FAD mutants alone with 4-week culture (Fig 4d). The accumulation of DNA damage could also contribute to cell death in AD (58).

Here we checked γ-H2AX expression as a marker for DNA damage. Ectopic expression of progerin could significantly upregulated γ-H2AX expression in both mAP group and mcherry control group in 4 weeks, whereas the difference between mcherry control group and mAP group was barely observed (Fig S6a,b). One possible explanation is that cells with FAD mutants alone might not be aged enough to develop detectable DNA damage.

## Discussion

## AD can be considered a laminopathy

Starting from the nuclear lamina, *via* the trans-membrane LINC complex to the cytoskeleton filaments, the nucleoskeleton and the cytoskeleton form a network of physically interconnected cellular components (59). The nuclear lamina plays a vital role in the signal transmission between the extracellular environment, cytoplasm, and nucleus (60). During the aging process, age-related pathogenesis may occur due to changes in cellular mechanical properties resulting from disrupting the nucleocytoskeleton’s integrity. This, in turn, could lead to dysfunctional changes. Protein aggregation is a common feature in most neurodegenerative diseases and could be linked to the disturbance in cell mechanics that occurs during the aging process.

Laminopathies are mainly caused by mutations in the *LMNA* gene and manifest nuclear architecture disruption (61). One of the laminopathies is HGPS, a premature aging disease (5). Observations have revealed several similarities between premature aging diseases and physiological aging. These similarities include instability in both genomic and proteomic structures, an increase in oxidative stress, and impaired DNA repair mechanisms (62). Additionally, in a drosophila model of AD, it was indicated that disruption of lamin led to relaxation of heterochromatin, activation of the cell cycle, and ultimately, cell death, all of which contribute to neurodegeneration (63). Furthermore, numerous studies have demonstrated alterations in nuclear morphology and increased lamin A in individuals with Alzheimer’s disease (27–30). One group also observed reduced ZMPSTE24, a protein that plays a major role in the cleavage of the farnesylated tail in prelamin A, in patient’s brain (64). These findings suggested that farnesylated lamin A is involved in AD. Thus, Alzheimer’s disease could be viewed as a form of acquired laminopathy.

Our study revealed that overexpression of lamin A in ReN cells resulted in elevated oxidative stress, reactivation of the cell cycle, and ultimately cell death. The presence of progerin, in particular, exacerbated these phenotypes, highlighting the potential role of lamin A in the development of AD pathology. As oxidative stress, cell cycle re-entry, and cell death are all crucial events in the aging process, these results suggest that the expression of lamin A or progerin may create an aging microenvironment conducive to the development of disease.

### Current AD *in vivo* and *in vitro* models

Currently, most of the AD drug candidates (>99.6% since 2002) in clinical trials fail to demonstrate sufficient clinical efficacy (65). A significant challenge in Alzheimer’s disease research is the absence of standardized and effective models. There are three main types of experimental models available: animal models, human post-mortem tissues, and cell culture models. Animal models are typically time-consuming and expensive. The majority of AD animal models are transgenic mice, with over 200 models developed to date (according to Alzforum). However, most clinical trials have failed, despite the efficacy of the compounds used to slow the disease in mouse models (66, 67). The reasons for the lack of translational success in mouse models are complicated. Due to the sequence differences between mouse Aβ and human Aβ (34), most mouse models utilized one or multiple FAD mutations to introduce transgenic human Aβ. Although these FAD models can generate some AD features, none of them can recapitulate the complete disease profile. Most of them do not show either amyloid plaques or tau tangles (68). Mouse and human tau share only 88% sequence homology and endogenous mouse tau could inhibit the aggregation of human tau (33, 69). This could explain the lack of tau tangles in these transgenic mice. More recently, knock-in mice were developed, which were considered more physiologically relevant (70). However, the pathological phenotypes and disease progression are still inconsistent, and if these knock-in mouse models are representative still needs to be validated.

### Development of an accelerated AD in vitro model

Human tissue or cells can overcome the interspecies differences encountered in animal models. However, a significant limitation in creating representative patient-derived models is the insufficient availability of high-quality post-mortem tissue. Consequently, most human-based models rely on induced pluripotent stem cells (iPSCs) (35). Nonetheless, there are currently no standardized protocols for generating and maintaining these cell lines, and even in 3D cultures, it takes several months to observe AD phenotypes (38). Additionally, a concern with iPSC-derived models is that the physiological age of the iPSCs may be reset, while AD is a late-onset disease. Thus, generating aged cells to study AD has become a pressing issue. Increased lamin A in AD patients might retain the farnesylated tail, given the observation that ZMPSTE24 is downregulated in AD brains (64). And progerin is the permanently farnesylated lamin A because of the loss of the ZMPSTE24 cleavage site (5). Moreover, HGPS exhibits molecular characteristics similar to those of natural aging, making it an effective model for aging research. Thus, using progerin to disrupt nuclear architecture could serve as a valuable strategy for emulating an aging environment. In one study, progerin was shown to induce age-related phenotypes in iPSC-derived neurons, successfully reproducing disease-specific phenotypes in Parkinson’s Disease (PD) (71).

Here we have put forward the idea that lamin A or progerin expression could create an environment conducive to the development of AD pathology. To test this hypothesis, we introduced exogenous lamin A or progerin into ReN cells containing FAD mutations to determine if AD-related features could be amplified. Our results demonstrated that ReN cells containing both progerin and FAD mutations exhibited a significant increase in the Aβ42/Aβ40 ratio and tau phosphorylation within just 3-4 weeks (Fig 3). Additionally, Aβ aggregation was checked, and we observed stronger Aβ oligomer staining (Fig S3e) and more Aβ fibril formation (Fig 3e) in these cells after a 4-week culture period. Furthermore, we noted a significant increase in cell cycle re-entry and cell death after progerin expression (Fig 4), which are crucial events in neurodegeneration. We examined cell-cycle-related regulators and observed that progerin expression could significantly increase the mRNA level of p16 and cdk4/6 (Fig S5). These findings suggest that progerin expression could accelerate the progression of AD-related phenotypes.

We hypothesize that cells containing only FAD mutations are not typically subject to aging in most circumstances. Therefore, their cell mechanics remain intact so they can clear toxic proteins. However, the balance of the nucleoskeleton is disrupted and the cellular environment becomes stiffer after progerin overexpression, making cells more vulnerable and leading to increased protein aggregation, cell cycle re-entry, and cell death (Fig 5a).

**Figure 5.**
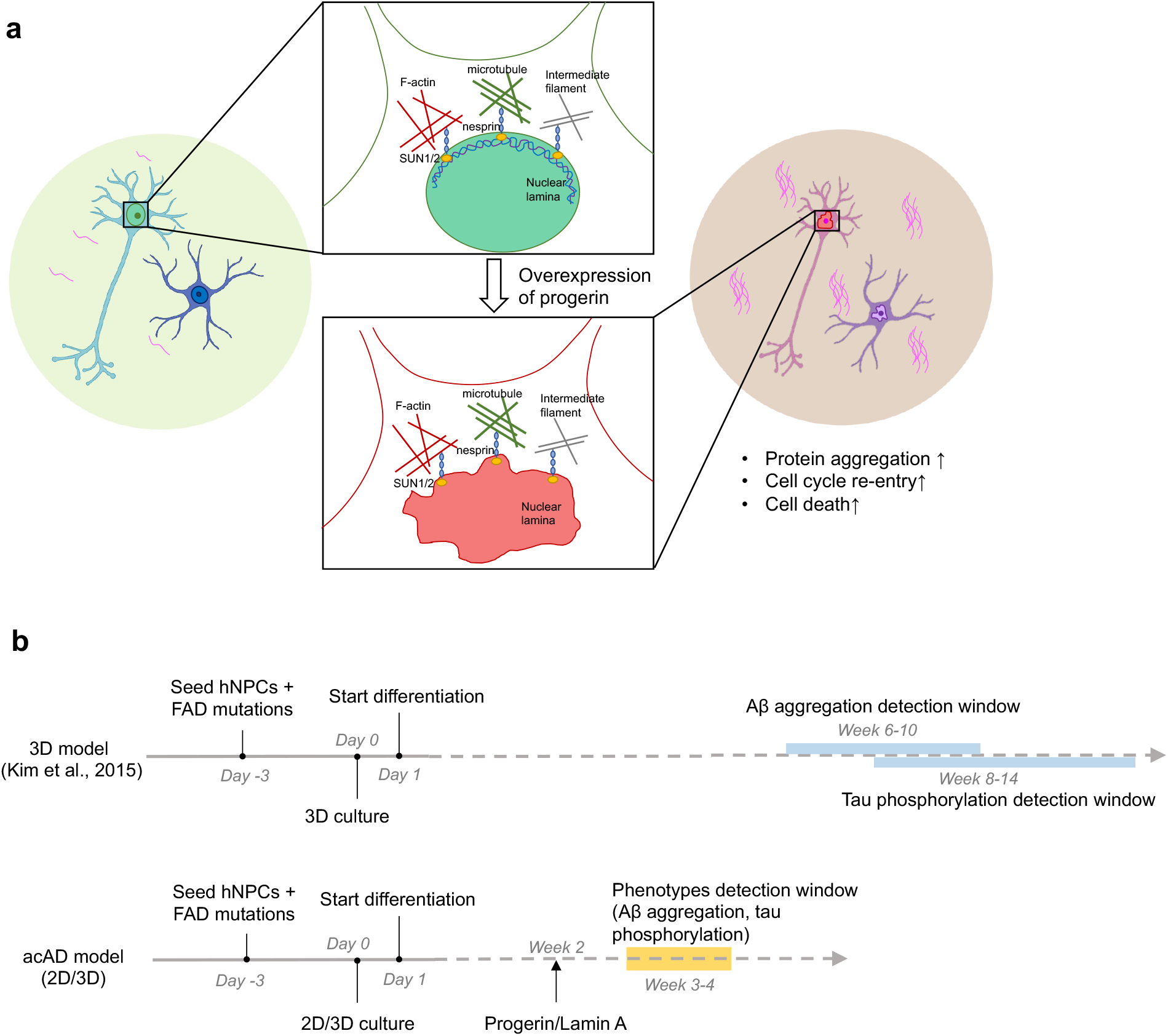
Proposed model of accelerated aging with progerin intervention. (a). We hypothesized that progerin could provide an aging environment for neurodegeneration. Cells containing only FAD mutations are not typically subject to aging in most circumstances. The connection between nucleoskeleton and cytoskeleton is integrated, and cell mechanic is well maintained. After progerin expression, the balance of the nucleoskeleton is disrupted and the cell environment could be stiffer, which makes cells vulnerable and results in more protein aggregation, cell cycle re-entry and cell death. (b). Comparison of schematic timeline between acAD model and a 3D model from Kim et al., 2015. Neural progenitor cells are differentiated from Week 0 in both protocols. Amyloid plaques and phosphorylated tau, two important AD hallmarks, were observed in the acAD model after 4-week differentiation, while it takes much longer in a well-characterized 3D AD model.

Compared to the time-consuming animal models and rejuvenated iPSC-derived models, our system served as a more efficient model. This system was built on the well-characterized, leading 3D AD cellular model (38), and therefore we adapted their timeline for comparison (Fig 5b). Our model demonstrated an accelerated manifestation of both increased tau phosphorylation and fibril formation within a significantly shorter timeframe of 4 weeks, in contrast to the traditional AD cellular models which typically require several months of experimental time. Meanwhile, traditional monolayer cell culture models fail to mimic the true brain architecture, resulting in Aβ diffusion in the culture medium (72). Our system can be utilized in both 2D and 3D cultures, providing the ability to easily adjust cell density and extracellular matrix thickness for a variety of analyses. Consequently, this accelerated AD model may be a more feasible platform for drug screening and investigating AD mechanisms, including neuronal death. Our future work will compare the transcriptomic and proteomic profiles among different groups. These profiles will be used to further validate our model, aid us in exploring potential new pathways involved in AD, and identify early biomarkers of the disease.

To summarize, our study has established a link between lamin A and AD pathology. We have demonstrated that by inducing progerin expression in FAD-mutant cells, we can create an aged environment and generate strong AD features in a short amount of time (Fig 5). This accelerated AD model could prove to be an effective tool for AD research. Furthermore, this progerin-induced aging can be used as a general approach for modeling other late-onset diseases.

## 4. Materials and Methods

### 4.1 hNPC cell culture

ReNcell VM immortalized human neural progenitor cells (hNPC) were purchased from EMD Millipore, initially derived from the ventral mesencephalon region of human fetal brains. Cells were expanded in cell culture plates coated with Matrigel (Corning) and grown in BrainPhys Neuronal Medium (STEMCELL) containing 20 ng/ml bFGF (R&D Systems), 20 ng/ml EGF (Millipore Sigma), 10U/ml heparin (Sigma-Aldrich), B27 supplement (Thermo Fisher). For differentiation, cells were cultured in described medium lacking bFGF and EGF. For 3D cell culture on 4-chamber slides, 50ul cold Matrigel was added to 50ul cell suspension on ice. Matrigel/cell mixture was further diluted by adding 400ul of the cold ReN differentiation medium and then seeded on chamber slides. After overnight 37°C incubation, 200ul of prewarmed ReN differentiation medium was added to each chamber.

### 4.2 Lentivirus packaging and transduction

HEK293T cells were co-transfected with lentiviral plasmids and two virus packaging vectors, psPAX2 and pMD2.G (Addgene), utilizing Fugene 6 (Promega). Culture supernatants were collected at 48h and 72h post-transfection, and filtered through 0.45μm filters to remove any nonadherent 293T cells, then stored at −80°C. Next, ReN cells were infected by lentiviruses in media supplemented with Polybrene (Santa Cruz Biotechnology) with the final concentration of 8 μg/ml. The medium was changed every other day post-infection until the cells were harvested.

### 4.3 RNA isolation and quantitative PCR

Total genomic RNA was extracted with Trizol (Life Technologies, Carlsbad, CA, USA) and purified using the RNeasy Mini kit (Qiagen, Hilden, Germany) as per the manufacturer’s instructions. The RNA yield was determined by the NanoDrop 2000 spectrophotometer (Thermo Fisher Scientific, Waltham, MA, USA). 600ng of total RNA was converted to cDNA using the iScript Select cDNA Synthesis kit (Bio-Rad). Quantitative RT-PCR was performed in triplicates using SYBR Green Supermix (Bio-Rad) on the CFX96 Real-Time PCR Detection System (C1000 Thermal Cycler, Bio-Rad). All primers used in this study are listed in Table 1.

### 4.4 Genomic DNA extraction and quantitative PCR telomere assay

DNA samples were extracted from ReN cells with PureLink™ Genomic DNA Mini Kit (Invitrogen). The mean telomere length was assessed by the modified monochrome multiplex quantitative polymerase chain reaction method (1). Relative telomere length is shown as T/S ratio, which stands for the ratio of telomere repeat copy number to single copy gene copy number. All primers used in this study are listed in Table 1.

### 4.5 Western blot

Whole-cell lysates for immunoblotting were prepared by dissolving cells in Laemmli Sample Buffer containing 5% 2-mercaptoethanol (Bio-Rad). Protein samples were loaded on 4-15% polyacrylamide gels (BioRad) and transferred onto 0.45 µm pore-size nitrocellulose membranes (Bio-Rad) using the Turboblot (BioRad). After that, blots were blocked with 5% milk for 1h at room temperature. For phospho-tau, 5% BSA in TBS was used for blocking. Blots were incubated overnight at 4°C with primary antibodies. And then blots were probed with secondary antibodies for 1h at room temperature before ECL development and imaging (Bio-Rad). The primary antibodies used for immunoblotting are as follows: Lamin A/C antibody, (Abcam, 1:750); Lamin B1 antibody (Santa Cruz, 1:200); APP antibody (BioLegend, 1:400), total tau antibody (Santa Cruz, 1:200), PHF6 p-tau antibody (Santa Cruz, 1:200), γ-H2AX antibody (Abcam, 1:3000), and β-actin (1:5000, Sigma-Aldrich).

### 4.6 Cell cycle assay

Cells were harvested with accutase and then washed with PBS. Next, ice-cold 70% ethanol was added to the cells and then cells were incubated at 4°C for 1h. After PBS wash, samples were treated with RNase at 37°C for 30min to remove RNA content. 5μg Propidium Iodide (Invitrogen) was added to the samples for another 30 min incubation at 37°C. Flow cytometry was performed with FACS CantoII (BD), and the data were analyzed by FlowJo software.

### 4.7 Cell death assay

PI-annexin V apoptosis assay was performed according to the manufacturer’s instruction (Thermo Fisher, A35122). In brief, cells were harvested and rinsed with PBS and then resuspended and stained with 100 μL of 1x annexin V binding buffer, containing 5 μL of annexin V and 5 μL of PI, for 25 min in the dark at room temperature. Stained samples were analyzed by FACS CantoII (BD), and the data were processed by FlowJo software.

### 4.8 Oxidative stress assay

Cellular ROS Assay kit (Abcam, ab186027) was used to check the oxidative stress according to the manufacturer’s protocol. Cells were dissociated by accutase digestion, rinsed with PBS, and then incubated in 1x ROS Red Stock Solution for 30min at 37°C. Flow cytometry was performed with FACS CantoII (BD), and the data were analyzed by FlowJo software.

### 4.9 ELISA

Aβ40 and Aβ42 levels were mainly measured by Invitrogen amyloid-β human ELISA Kit (Thermo Fisher, KHB3481 and KHB3441) as per the manufacturer’s protocol. The conditioned media from ReN cells were collected and diluted by 1:3 or 1:9 with a dilution buffer provided by the manufacturer. A plate reader (Thermo Scientific) was used to quantify Aβ40 and Aβ42 ELISA signals.

### 4.10 Immunofluorescence staining

Cells were washed twice with PBS and then fixed in 4% paraformaldehyde (PFA) for 15 min at room temperature. After that, cells were permeabilized with 0.5% triton in PBS for 5 min at room temperature and then washed twice in TBS. Following blocking was done with 4% BSA in TBS for 1h at room temperature. Samples were then incubated with primary antibodies in 4% BSA in TBS overnight at 4°C. Primary antibodies were rinsed off with 5 washes of TBS. Samples were incubated with secondary antibodies in 4% BSA in TBS for 1h at room temperature protected from light before being washed 5 times in TBS. Primary antibodies used include: MAP2 antibody (Abcam, 1:1000); β-tubIII antibody (Abcam, 1:1000); GFAP antibody (Cell Signaling, 1:1000); Lamin A/C antibody, (Abcam, 1:500); Lamin B1 antibody (Santa Cruz, 1:200). Secondary antibodies include: Alexa Fluor 488 donkey anti-rabbit IgG (1:1000, Invitrogen), Alexa Fluor 594 donkey anti-rabbit IgG (1:1000, Invitrogen), Alexa Fluor 488 donkey anti-mouse IgG (1:1000, Invitrogen), Alexa Fluor 594 donkey anti-mouse IgG (1:1000, Invitrogen) and Alexa Fluor 644 donkey anti-rabbit IgG (1:1000, Invitrogen).

For BrdU staining, cells were incubated in a 10 uM BrdU (BD #550891) labeling medium for 12h. Cells were then washed with PBS, fixed and permeabilized. Afterward, cells were denatured in 2N HCl for 40 min at room temperature. Next, samples were incubated in Alexa Fluor 647 anti-BrdU antibody solution (1:1000, Invitrogen #B35133) at 4°C overnight. Samples were protected and stained in vectashield mounting medium with DAPI (Vector) sealed with a coverslip and stored in the dark. Fluorescence images were acquired with a Zeiss LSM 710 confocal microscope (Zeiss International, Oberkochen, Germany).

### 4.11 Amylo-glo staining

Cells were washed three times with 0.9% (wt/vol) NaCl solution. Following adding 100 µl of 0.05× Amylo-Glo working solution, 3D culture cells were incubated for 5 min at room temperature. And then, the staining solution was removed. 200 µl of 0.9% saline was added and followed with 5-min incubation.

Samples were washed three times with ddH2O, and further washed with 0.9% (wt/vol) NaCl solution three times. Samples were protected in antifade vectashield mounting medium without DAPI (Vector) and stored in the dark. Fluorescence images were acquired with a Zeiss LSM 710 confocal microscope (Zeiss International, Oberkochen, Germany).

### 4.12 Calcium preparation and imaging

Intracellular calcium labeling was prepared using Fluo-4 AM (Thermo Fisher Scientific) following manufacturer-provided protocols. Briefly, a 1mM stock solution was prepared in anhydrous DMSO. Cells were incubated in standard cell specific medium for 1 hour at 37°C with 1uM Fluo-4 AM. After incubation cells were rinsed with DPBS and then the fresh, appropriate medium was added. Cell samples were then immediately used for imaging using 488 nm laser to excite the Fluo-4 AM dye.

### 4.13 Data analysis

Statistical analyses were performed using GraphPad Prism 7 software. Data were analyzed using unpaired Student’s *t*-test for two groups. One-way and two-way analysis of variance (ANOVA) followed by post hoc multiple comparisons were used to compare the means of three or more groups. All experiments were repeated at least three times, and the results are presented as the mean ± SD. A *p* value < 0.05 was considered significant. Asterisks indicate statistical difference as follows: n.s., not significant; *p < 0.05; **p < 0.01; ***p < 0.001; ****p < 0.0001.

## Data availability

This study includes no data deposited in external repositories.

## Acknowledgments

We thank the Imaging Core and FACS core at the University of Maryland for their technical support. We would like to thank all the members of the Cao lab for their helpful discussions and suggestions.

