## Supplementary materials for "Development of an Accelerated Cellular Model for Alzheimer’s Disease"

#### Figure S1.

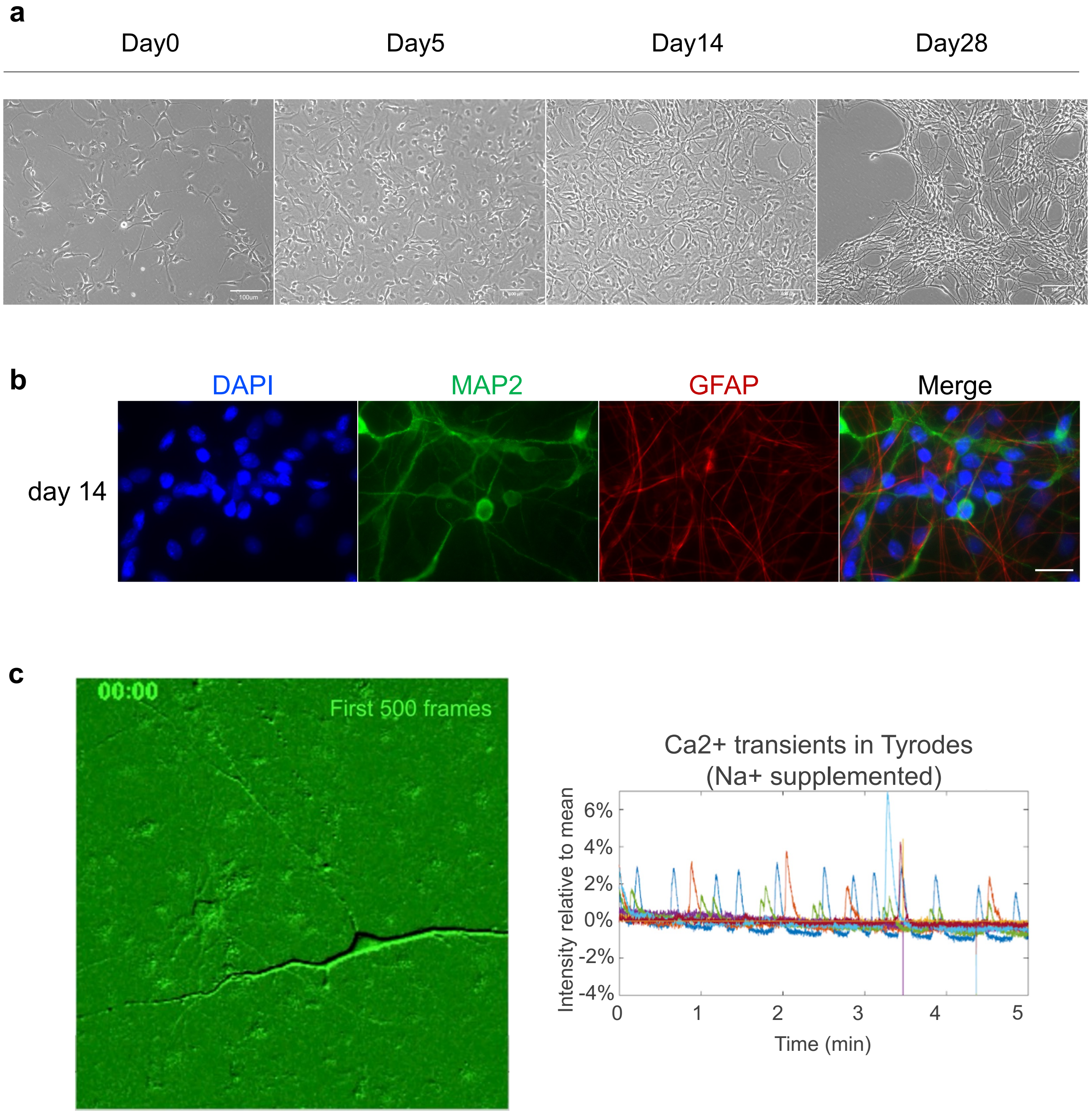


Figure S1. The process of ReN cell differentiation.
(a) ReN cells could grow neurites in two weeks. (Scale bar: 100um)
(b) Differentiated cells were positive with neuronal markers, MAP2 and astrocyte markers, GFAP. (Scale bar: 20um)
(c) Differentiated cells could generate Ca2+ transients in the Tyrode's solution supplemented with Na+.

**Figure S2.**

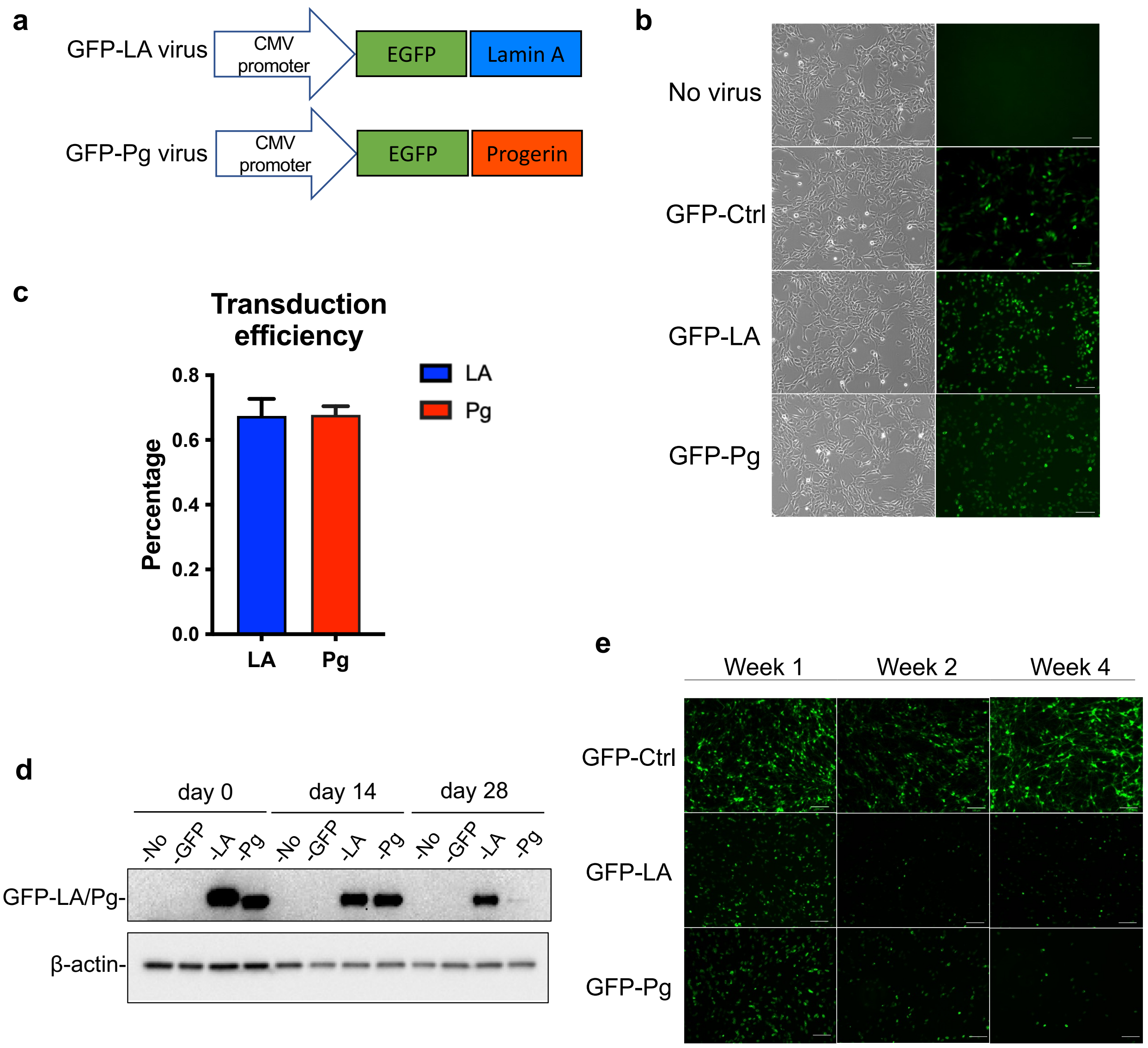


Figure S2. Lamin A- and progerin- transduction in ReN cells.
(a) Diagrams of lentiviral constructs. GFP-LA virus contains lamin A cDNA and GFP-Pg virus contains progerin cDNA. Both vectors contain EGFP signal.
(b)(c) Quantification of transduction efficiency. The transduction was successful, and the efficiency was 71.1% for GFP-LA virus and 64.8% for GFP-Pg virus after 2-day transduction.
(d) Western blot results of exogenous protein level and GFP signals during the differentiation. Both exogenous lamin A and progerin protein were decreased during the differentiation.
(e) GFP signal under the microscope. GFP signal was stable in GFP-ctrl cells, while GFP signal decreased during the differentiation in lamin A- or progerin- transduced cells. (Scale bar: 100um)

#### Figure S3.

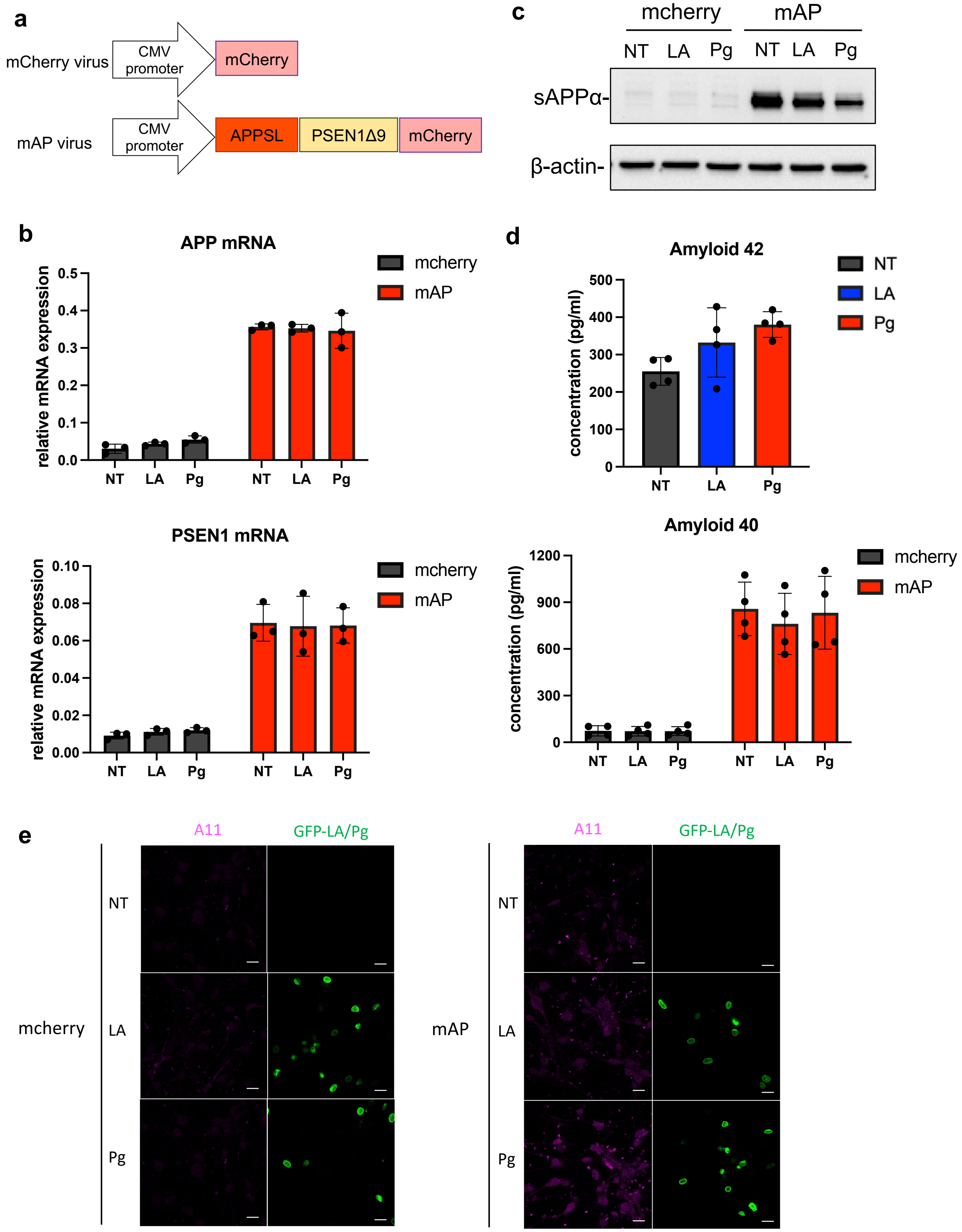


(Figure legend on the next page)

Figure S3. Combination of FAD mutations and progerin-transduction in ReN cells
(a) Diagrams of lentiviral constructs. A plasmid containing APP with both the K670N/M671L (Swedish) and V717I (London) mutations (APPSL) and PSEN1 with the Δ9 mutation (PSEN1(Δ9)) was a gift from Dr. Kim's lab.
(b) Quantification of mRNA level after FAD transduction. The transcription level of both APP and PSEN1 was upregulated after the transduction.
(c) Western blot of APP protein. sAPPα was increased in mAP group after the transduction.
(d) Protein concentration of Aβ40 and 42 in the culture medium after 3-week culture.
(e) Aβ oligomer staining after 3-week differentiation. Pink indicated Aβ oligomer staining, green indicates the GFP-tagged lamin A or progerin. (Scale bar: 20um)

### Figure S4.

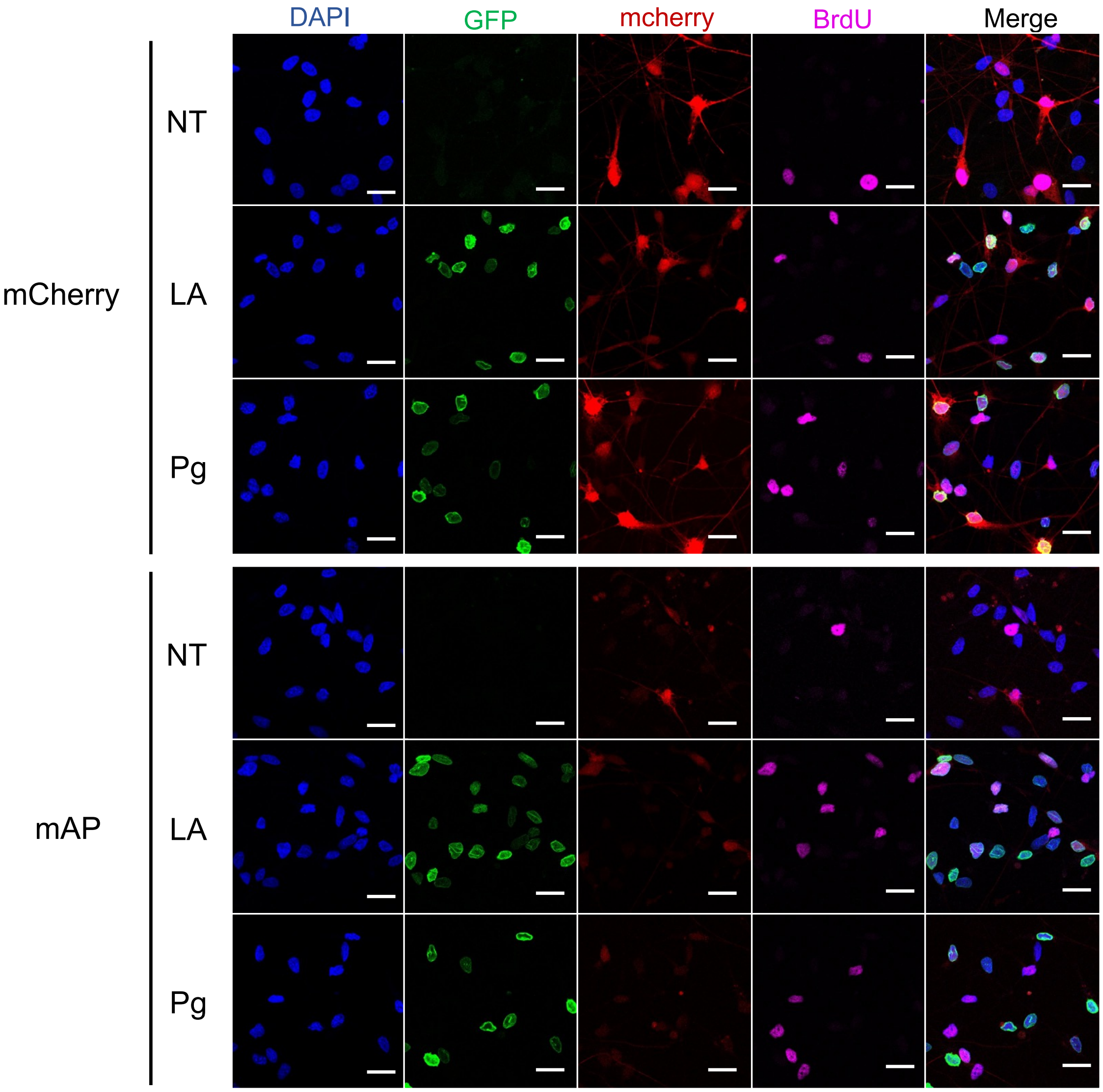


Figure S4. Immunofluorescence staining of BrdU in different groups after 4 weeks. Blue indicated the nucleus, green indicated the GFP-tagged lamin A or progerin, red indicated mcherry or mcherry-tagged APP and PSEN1, pink indicated BrdU staining. (Scale bar: 20um)

**Figure S5.**

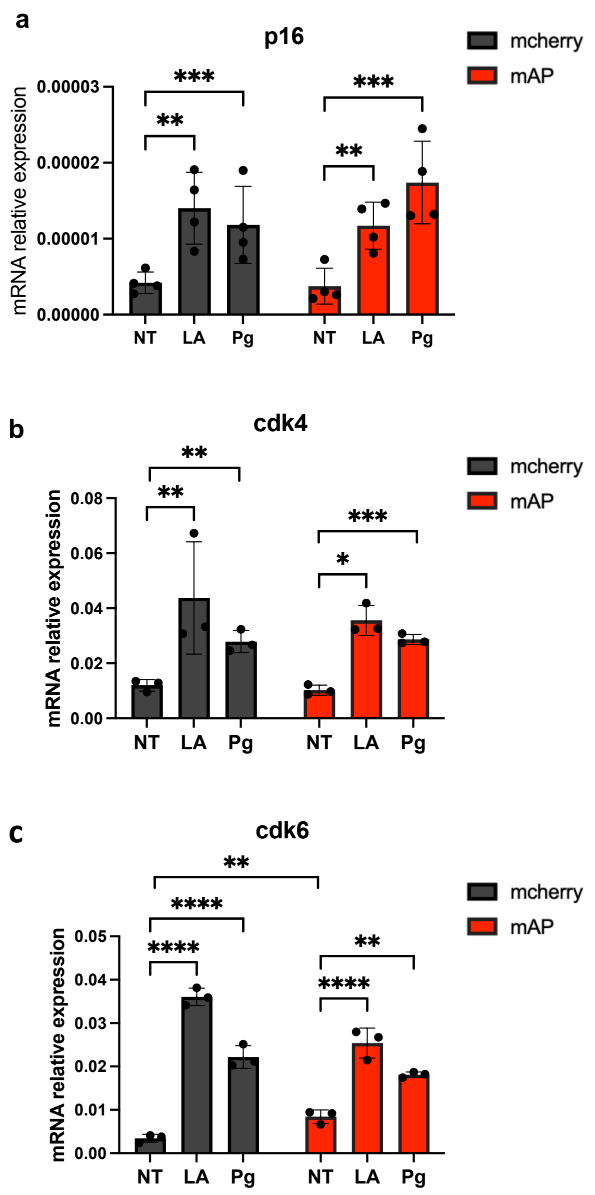


Figure S5. The mRNA expression of cell-cycle-related regulators after 4 weeks.
(a). Quantification of p16 mRNA expression. Within mcherry and mAP groups, p16 mRNA level was significantly increased after lamin A expression. Same trend was observed after progerin expression. Results were generated from four biological replicates. n.s., not significant; *$p < 0.05$; **$p < 0.01$; ***$p < 0.001$; ****$p < 0.0001$.
(b). Quantification of cdk4 mRNA expression. Within mcherry and mAP groups, cdk4 mRNA level was significantly increased after lamin A expression. Same trend was observed after progerin expression. Results were generated from three biological replicates. n.s., not significant; *$p < 0.05$; **$p < 0.01$; ***$p < 0.001$; ****$p < 0.0001$.
(c). Quantification of cdk6 mRNA expression. mRNA level of cdk6 was higher in the cells with FAD mutants alone than the cells with mcherry control alone. Within mcherry and mAP groups, cdk6 mRNA level was significantly increased after lamin A expression. Same trend was observed after progerin

expression. Results were generated from three biological replicates. n.s., not significant; *$p < 0.05$; **$p < 0.01$; ***$p < 0.001$; ****$p < 0.0001$.

**Figure S6.**

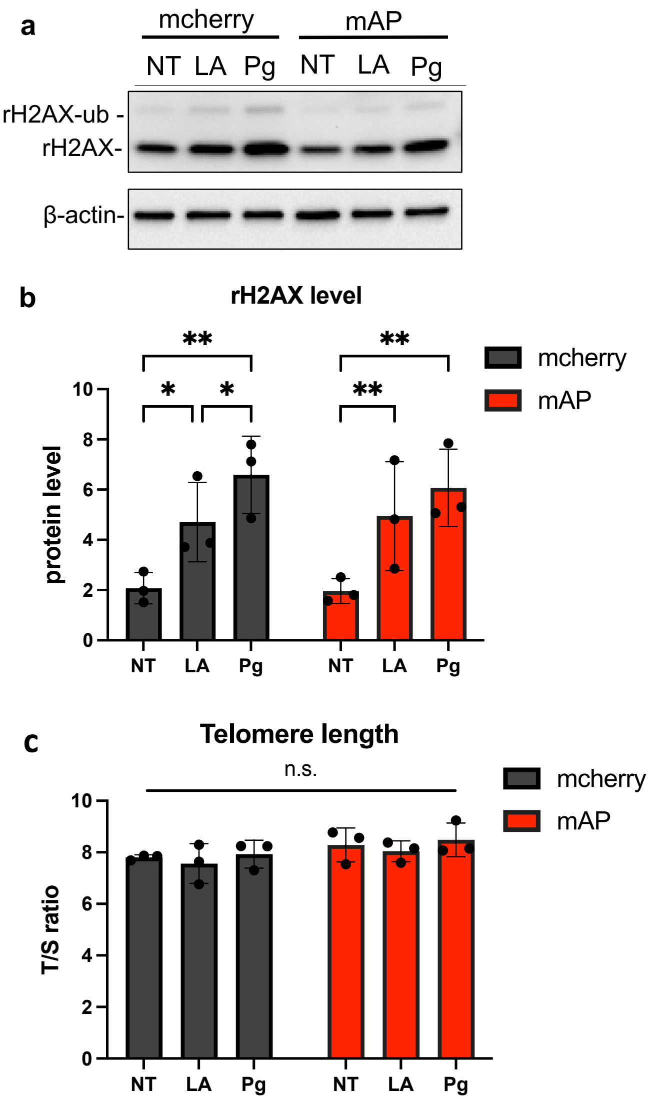


Figure S6. DNA damage and telomere length after progerin addition.
(a)(b). Western blot of γH2AX. γ-H2AX was upregulated after ectopic lamin A expression in both mcherry control group and mAP group after 4 weeks. Same trend was observed after progerin expression. Results were generated from three biological replicates. n.s., not significant; *$p < 0.05$; **$p < 0.01$; ***$p < 0.001$; ****$p < 0.0001$.
(c). Quantification of telomere length with qPCR after 4 weeks. Telomere length did not show significant change with FAD intervention or progerin intervention. Results were generated from three biological replicates. n.s., not significant.

| Gene | Forward (5'-3') | Reverse (5'-3') |
|---|---|---|
| **ACTB** | CTGGAACGGTGAAGGTGACA | AAGGGACTTCCTGTAACAATGCA |
| **Lamin A** | GCAACAAGTCCAATGAGGACCA | CATGATGCTGCAGTTCTGGGGGCTCTGGAT |
| **Lamin C** | CTCAGTGACTGTGGTTGAGGA | AGTGCAGGCTCGGCCTC |
| **Lamin B1** | GGAGAATCGTTGTCAGAGCCTT | TGCGGCTTTCCATCAGTTCT |
| **APP** | TGGGTTCAAACAAAGGTGCA | GTTCTGCTGCATCTTGGACA |
| **PSEN1** | GCAGTATCCTCGCTGGTGAAGA | CAGGCTATGGTTGTGTTCCAGTC |
| **P16** | GAGCAGCATGGAGCCTTC | CGTAACTATTCGGTGCGTTG |
| **Cdk4** | CCATCAGCACAGTTCGTGAGGT | TCAGTTCGGGATGTGGCACAGA |
| **Cdk6** | CCAGGCAGGCTTTTCATTCA | AAGTATGGGTGAGACAGGGC |
| **Telomere** | CGGTTTGTTTGGGTTTGGGTTTGGGTTTGGGTTTGGGTT | GGCTTGCCTTACCCTTACCCTTACCCTTACCCTTACCCT |
| **β-globin** | GCTTCTGACACAACTGTGTTCACTAGC | CACCAACTTCATCCACGTTCACC |

Table 1. qPCR and telomere length assay primer list
